# Capillary regression leads to sustained local hypoperfusion by inducing constriction of upstream transitional vessels

**DOI:** 10.1101/2023.10.28.564529

**Authors:** Stephanie K. Bonney, Cara D. Nielson, Maria J. Sosa, Andy Y. Shih

## Abstract

In the brain, a microvascular sensory web coordinates oxygen delivery to regions of neuronal activity. This involves a dense network of capillaries that send conductive signals upstream to feeding arterioles to promote vasodilation and blood flow. Although this process is critical to the metabolic supply of healthy brain tissue, it may also be a point of vulnerability in disease. Deterioration of capillary networks is a hallmark of many neurological disorders and how this web is engaged during vascular damage remains unknown. We performed *in vivo* two-photon microscopy on young adult mural cell reporter mice and induced focal capillary injuries using precise two-photon laser irradiation of single capillaries. We found that ∼63% of the injuries resulted in regression of the capillary segment 7-14 days following injury, and the remaining repaired to re-establish blood flow within 7 days. Injuries that resulted in capillary regression induced sustained vasoconstriction in the upstream arteriole-capillary transition (ACT) zone at least 21 days post-injury in both awake and anesthetized mice. This abnormal vasoconstriction involved attenuation of vasomotor dynamics and uncoupling from mural cell calcium signaling following capillary regression. Consequently, blood flow was reduced in the ACT zone and in secondary, uninjured downstream capillaries. These findings demonstrate how capillary injury and regression, as often seen in age-related neurological disease, can impair the microvascular sensory web and contribute to cerebral hypoperfusion.

**SIGNIFICANCE:** Deterioration of the capillary network is a characteristic of many neurological diseases and can exacerbate neuronal dysfunction and degeneration due to poor blood perfusion. Here we show that focal capillary injuries can induce vessel regression and elicit sustained vasoconstriction in upstream transitional vessels that branch from cortical penetrating arterioles. This reduces blood flow to broader, uninjured regions of the same microvascular network. These findings suggest that widespread and cumulative damage to brain capillaries in neurological disease may broadly affect blood supply and contribute to hypoperfusion through their remote actions.

## INTRODUCTION

Vascular contributions to cognitive impairment and dementia (VCID) include small, but wide-spread ischemic and hemorrhagic injuries called microinfarctions and microbleeds, respectively^1,2^. These lesions are <3 mm in size and may only occupy 1-2 mL of the total brain volume. However, their remote effects can depress neural function and contribute more broadly to brain dysfunction and cognitive decline^3^. While these micro-lesions are well-known features of VCID and believed to result largely from pathologies of brain arterioles, even smaller disruptions occur in capillaries, the finest and most delicate vessels in the brain. Capillaries are the site of amyloid beta buildup during Type 1 cerebral amyloid angiopathy, a form of small vessel disease^4^, and reactive oxygen species induced by amyloid beta toxicity can cause damage to the capillary endothelium and neighboring pericytes^5^. Microbleeds can occur in the absence of capillary amyloid and may result from proteolytic degradation of the endothelium by pericytes or trapped neutrophils^6-8^. These events contribute to capillary network rarefaction, as commonly seen in aging and VCID^9,10^, and evade detection by clinical imaging due to their exceedingly small size. Yet, they likely contribute to disease progression in covert and insidious ways. Since the capillary network is dense and redundant in its connectivity^11,12^, it is unclear whether focal capillary changes could lead to broader, yet unrecognized impairments of brain perfusion.

Capillaries sense neuronal activity via Kir2.1 and TRPA1/Panx1 channels on the endothelium^13,14^ and Kir2.2 on pericytes^15^. This initiates a capillary-to-arteriole conductive hyperpolarization that is propagated to upstream arteriole-capillary transitional (ACT) vessels and penetrating arterioles leading to reduced intracellular calcium (Ca^2+^), mural cell relaxation, and increased blood flow back downstream into the capillary networks^16^. The conductive properties of the microvasculature are also dependent upon endothelial and pericyte gap junctions^17^. The ACT zone is where many vasomodulating signals converge, making it an important control point for blood flow^18-20^. Vessel segments in this region are surrounded by α-smooth muscle actin-expressing ensheathing pericytes, and may also include precapillary sphincters at upstream branch points. The ACT zone is highly sensitive to brain activity, and is the first to dilate during neurovascular coupling^21,22^. However, it is also susceptible to pathology and exhibits sustained constriction after transient cerebral ischemia^23^. Critically, depolarizing electrical stimulation of capillary pericytes leads to a conductive wave of Ca^2+^ increase upstream that elicits robust vasoconstriction in ACT vessels^17^. These findings cast the capillary network as a “sensory web” that responds to physiological activity, but potentially also to pathophysiological stimuli.

To test this possibility, we used *in vivo* two-photon imaging in the mouse cerebral cortex to induce precise injuries, and then tracked the consequence of these injuries over weeks in both anesthetized or awake mice. We asked: (1) How does capillary injury affect the flow and dynamics of upstream ACT and penetrating arteriole zones? (2) Are there lasting effects of capillary injury on blood flow?

## RESULTS

### Focal capillary injury induces vessel regression

To understand how capillary injuries affect the microvascular network, we performed *in vivo* two-photon imaging on mural cell reporter mice (PdgfrβCre-tdTomato)^24^ and used a modified model of focal capillary injury involving rupture of the vascular wall^25^. Capillaries were identified by distinguishing thin-strand and mesh pericytes on capillary vessels from morphologically distinguished ensheathing pericytes on ACT vessels^18^. In general, ruptured capillary segments were 5 to 8 branch orders from penetrating arterioles. Injuries were induced by focusing <3 μm diameter circular laser line-scan (∼100-154 mW at 800 nm)(**Supp. Fig. 1A**) directly on the vessel lumen as visualized by the perfusion of intravenous (i.v.) dye (70kDa FITC-dextran) (**Fig. 1A**). Laser power was applied in 20 second (s) increments for a total of 20-80 s until there was indication of halting blood flow and dye leakage (**Fig. 1A, B**). Sham injuries were performed in separate vascular networks using identical laser powers (**Supp. Fig. 1A**) localized next to similarly sized capillary segments (**Supp. Fig. 1B**) without damaging the vessel (**Fig. 1B & Supp. Fig. 1C**). Capillary diameter did not influence the laser power or time needed to rupture the vessel (**Supp. Fig. 1D, E**). Although, an increase in laser power and not time was needed for injuries deeper into the cortex (**Supp. Fig. 1F, G**).

**Figure 1.**
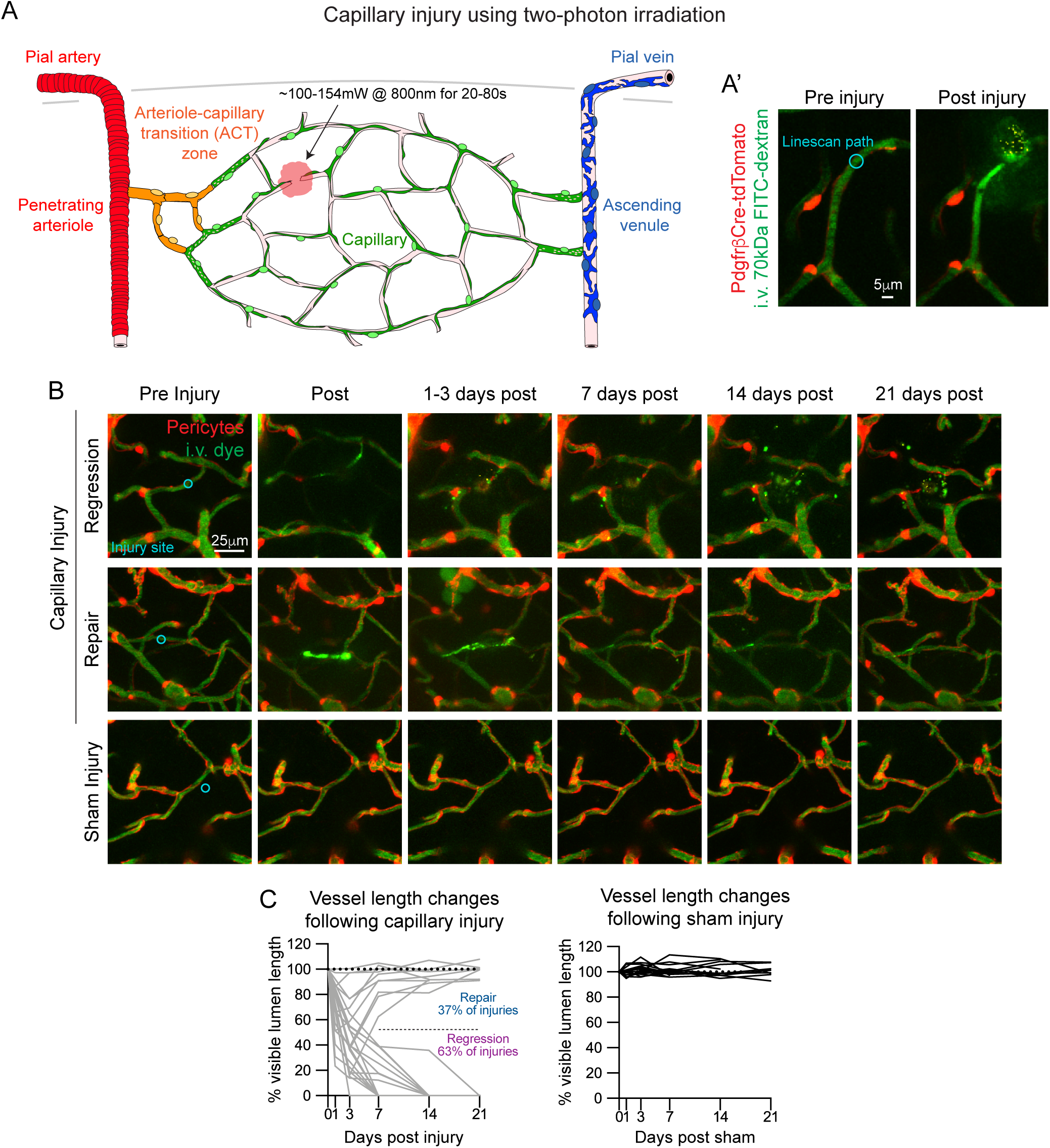
Focal capillary injury induced by optical laser ablation. **(A)** Schematic of a cortical microvascular network with location of two-photon laser-induced capillary injury. Microvascular zones are highlighted including the pial artery and penetrating arteriole (red), arteriole-capillary transition zone (orange), capillary zone (green), ascending venule and pial vein (blue). **(A’)** Representative *in vivo* image of a capillary injury in a PdgfrβCre-tdTomato mouse pre-injury with line-scan path (cyan) and ∼10 minutes post-injury. Pericytes are shown in red and i.v. dye (70kDa FITC-Dextran) labeling vessels in green. **(B)** Representative *in vivo* images of a capillary regression, repair, and sham event pre- and post-injury (∼10 minutes), and then at various days post-injury. **(C)** Graphs of percent change in length of the injured (gray; left) or sham (black; right) capillaries based on visible i.v. dye in the lumen pre-injury (0 day) and various days post-injury. Overall, 10/27 injuries (37%) resulted in capillary repairs and 17/27 (63%) resulting in regressions, in experiments conducted over 14 mice. Sham injuries = 15 conducted over 12 mice.

To understand the consequence of focal capillary injury, injured regions were imaged longitudinally for 21 days. One to three days post-injury (dpi), the majority of vessels remained disconnected and lacked blood flow (**Fig. 1B & Supp. Fig. 1H**). Quantification of vessel length demonstrated that most vessels (63% of injuries) had completely receded back to their branch points 7 to 14 dpi (**Fig. 1C**). We defined these as regression events. A smaller portion of injured vessels (37%) reconnected and re-established blood flow 7 dpi (**Fig. 1C & Supp. Fig. 1H**). We called these repair events. Sham injuries did not change the vessel segment length. The occurrence of regression or repair events was not due to differences in laser power, time, diameter or vessel length (**Supp. Fig. 2A-D**). No consistent trend in capillary response was seen with vascular branch order (**Supp. Fig. 2E**), but superficial capillaries (0-50 μm cortical depth) were more likely to regress (**Supp. Fig. 2F**). Overall, these findings show that precise laser line-scans can be used to induce focal capillary injury *in vivo*, with roughly ⅔ of injured vessels experiencing regression and ⅓ undergoing gradual repair.

### Pericyte death may be induced by capillary injury, but remodeling of neighboring pericytes ensures coverage

Following capillary injuries, death of nearby pericytes was occasionally observed 3 dpi (22% of injuries), leaving a portion of the capillary network transiently uncovered by pericytes (**Supp. Fig. 3A**). Pericyte loss was due to direct injury to a cellular process with the line-scan, or indirectly from the resultant bleed. The somata of dying pericytes were often quite distant from the injury site, highlighting the vulnerability of these cells to injury of their processes or perturbations in their microenvironment. In contrast, pericyte loss was not observed following sham injuries, suggesting that local thermal damage to the parenchymal tissue does not drive pericyte loss.

Pericyte death triggered remodeling of neighboring pericytes, which extended their processes into areas of the endothelium lacking pericyte coverage (**Supp. Fig. 3B**)^26,27^. Pericyte process growth was evident at 3 dpi, and in most cases, remodeling was complete by 14 dpi. In sham injuries, pericyte process territory negotiation was observed, evidenced by small local fluctuations in process length, which are known to occur in the healthy mouse brain^26,27^ (**Supp. Fig. 3C**). Remodeling pericyte processes achieved an average extension of 62.67 μm, compared to 8.93 μm for pericytes at sham sites (**Supp. Fig. 3D**). Analysis of uncovered vascular length following pericyte death confirmed that most vessels lacking pericyte contact were recovered by 7-14 dpi (**Supp. Fig. 3E**).

As shown in past studies involving direct ablation of pericytes via their somata^26,27^, loss of pericyte coverage with capillary injury resulted in dilation of capillaries (**Supp. Fig. 3F**). On average, uncovered vessels dilated to 117.5% of baseline, and returned approximately back to their basal diameter following recoverage by pericyte remodeling (**Supp. Fig. 3G**). No significant vessel diameter changes were observed at sham injury sites. Thus, laser-induced capillary injury leads to loss of pericyte coverage in approximately ⅕ of cases, but remodeling of neighboring cells ensures that pericyte coverage is regained over days/weeks.

### Focal capillary injury induces a local microglia response

To assess the focality of the capillary injuries, we utilized PdgfrβCre-tdTomato; Cx3cr1-GFP mice to observe microglial activity following injury. We quantified the area of GFP-positive fluorescence within a 25 μm radius surrounding the injury site, which encompassed the size of the initial microglia response to capillary injury. We found that microglia rapidly migrated to the site 1 dpi (**Fig. 2A, B**). In regression events, microglia occupancy at the injury site remained significantly high 7 and 14 dpi. In contrast, microglial presence at the injury site decreased to baseline 14 dpi in repair events. Sham injuries did not result in a robust or prolonged microglia response at the site of laser ablation (**Fig. 2A, B**). Outside of the irradiation site, we observed no change in microglia density following regression, repair, or sham events (**Fig. 2C**). This demonstrates that laser-induced capillary injuries initiate a highly focal inflammatory response that is prolonged in the event of vessel regression, but does not induce inflammatory conditions along other regions of the broader vascular network.

**Figure 2.**
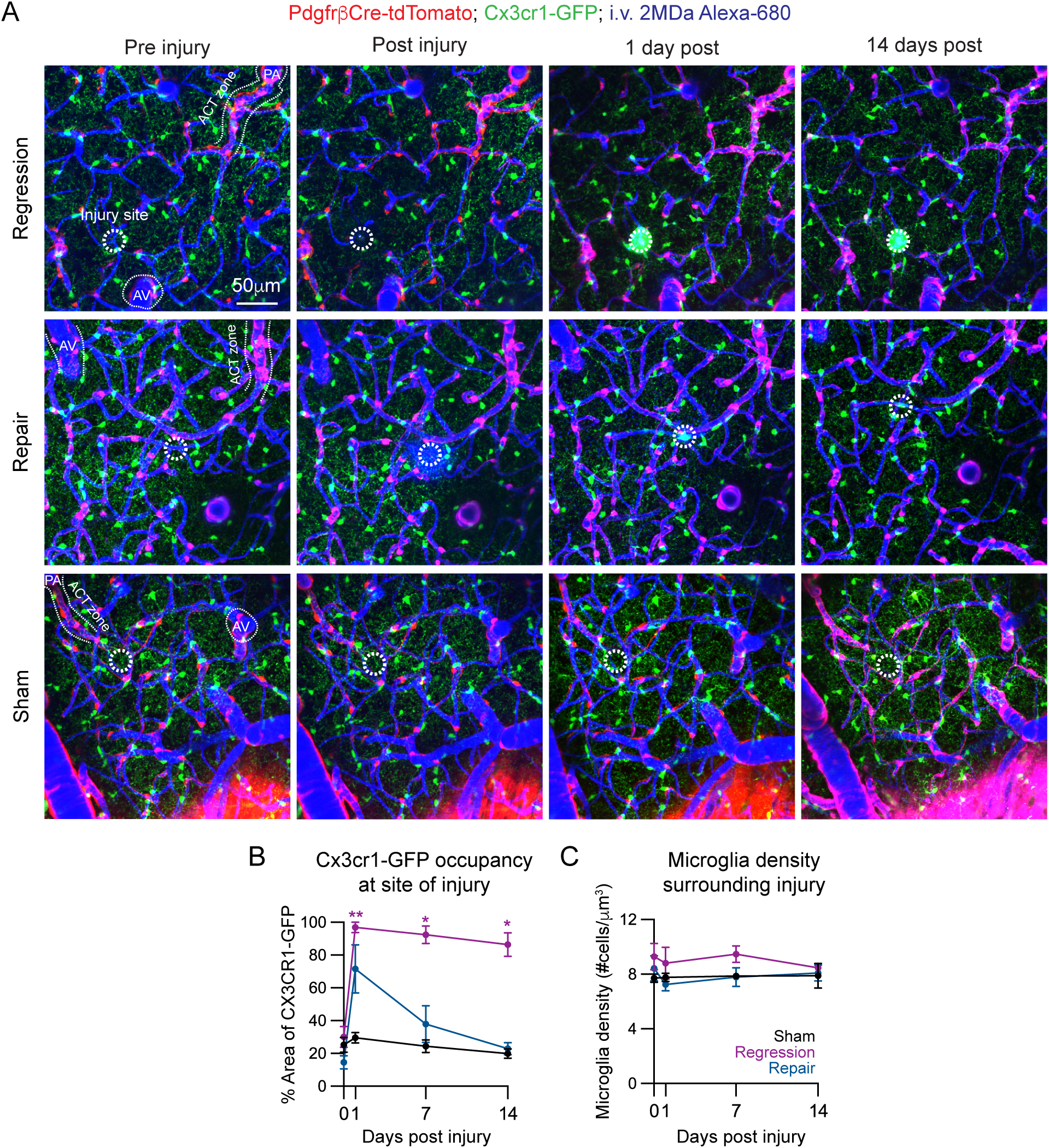
Microglia activation is concentrated only around injury site. **(A)** Representative *in vivo* images of a capillary injury in a PdgfrβCre-tdTomato; Cx3cr1-GFP mouse pre, post (∼10 minutes), 1-, and 14-days post-capillary injury. Pericytes are shown in red, microglia in green, and i.v. dye (2MDa Alexa 680-Dextran) labeling vessels depicted in blue. Microvascular zones such as penetrating arteriole (PA), arteriole-capillary transition (ACT) zone, and ascending venules (AVs) are shown to depict the highly focal nature of the laser-induced capillary injuries and resulting neuroinflammatory reaction (25 μm radius; white dashed circle). **(B)** Graph of percent area of Cx3cr1-GFP+ microglia at injury site pre-injury (0 day) and 1-, 7-, and 14-days post-injury. ANOVA followed by Dunnett’s multiple comparison test were performed: Regression event: 0 vs. 1 day **p=0.0074, 0 vs. 7 days: *p=0.0109, 0 vs 14 days: *p=0.0385. Sham injuries n=3, regression events n=3, repair events n=3; 2 mice. **(C)** Graph of Cx3cr1-GFP+ microglia density surrounding injury site (excludes injury site) pre-injury (0 day) and 1-, 7-, and 14-days post-injury. ANOVA tests were performed and no significant differences in microglia density were detected.

### Constriction of the arteriole-capillary transition zone following capillary regression

We next examined the effect of capillary injury on the broader microvascular network. For each experiment, we mapped the territory of morphologically distinct mural cells along the vascular tree to identify different vascular zones, including the penetrating arteriole, ACT zone, capillaries, and ascending venules (**Fig. 3A**). We then measured the resting diameter of each vessel segment to understand if focal capillary injury initiates a response in other vascular zones. We monitored diameter throughout the acute phase, when the injured vessel underwent regression or repair (3-7 dpi), and the chronic phase, when the regression or repair was complete (14-21 dpi). Strikingly, we found a specific decrease in vessel diameter in the upstream ACT zone following capillary regression during both the acute and chronic phases (**Fig. 3B, C**). Some alterations were noted within the capillary zone in the event of regression but did not maintain a consistent pattern. There were no significant differences in vessel diameter across all vascular zones following sham or repair events during acute and chronic phases (**Supp. Fig. 4**). Therefore, repair and sham events were pooled and compared to regression events.

**Figure 3.**
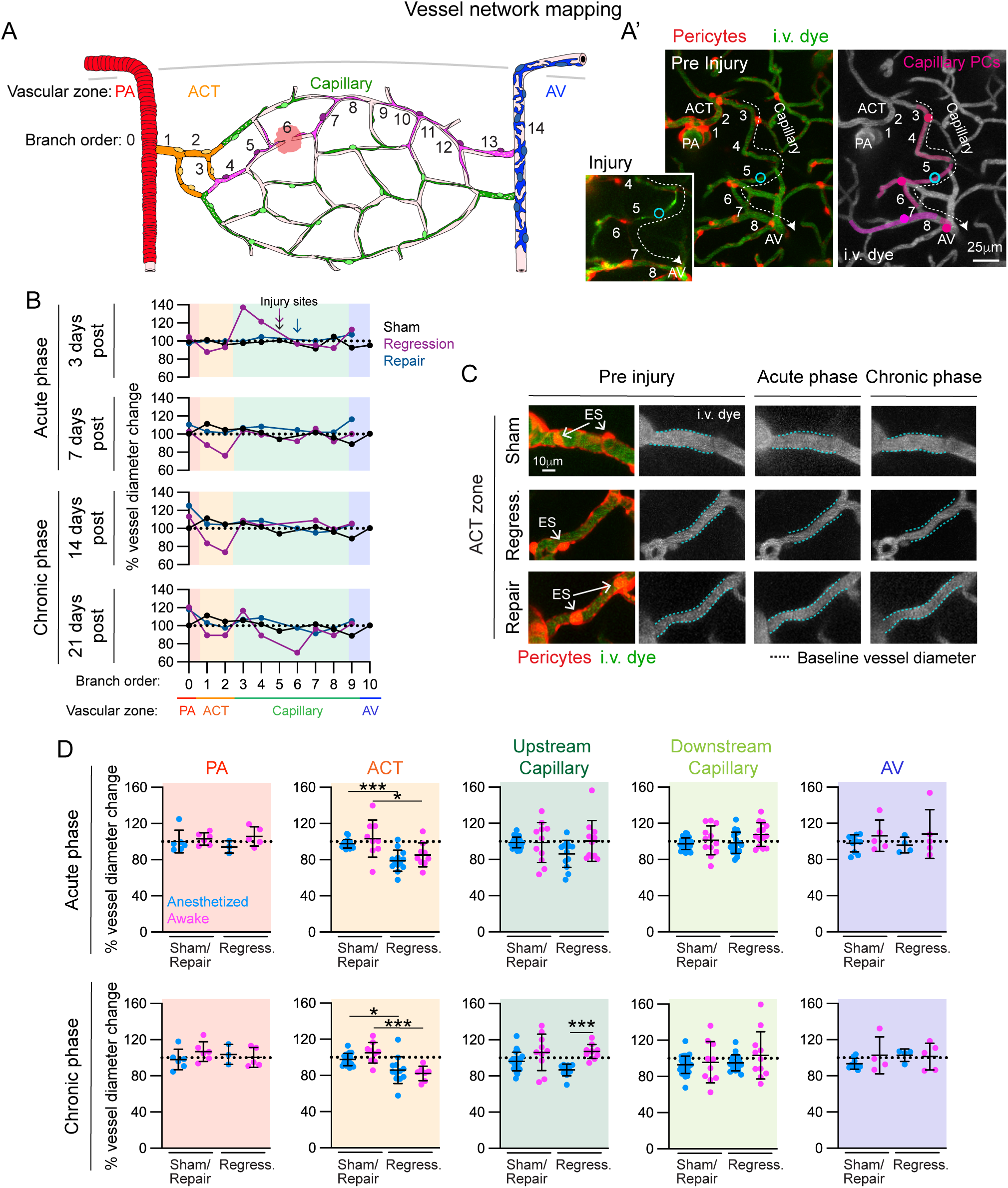
Chronic constriction of arteriole-capillary transition vessels occurs following laser-induced capillary regression. **(A)** Schematic of vessel network mapping by tracing pericyte territories (pink) and branch order from penetrating arteriole (PA) and arteriole-capillary transition (ACT) zone through the capillary network including the injury site to the ascending venule (AV). **(A’)** Representative *in vivo* image of a capillary injury with pericyte territories in a PdgfrβCre-tdTomato mouse pre-injury with the line-scan path (cyan) and ∼10 minutes post-injury (inset). Pericytes are shown in red and i.v. dye (70kDa FITC-Dextran) labeling vessels depicted in green. In the right panel, pericyte territories are shown in pink, as well as vascular branch order and blood flow direction (dash line and arrow). **(B)** Graph of percent change in vessel diameter from baseline across the microvascular zones (PA, ACT, Capillary, AV) in a vessel network of a sham (black), regression (purple), and repair (blue) event. Changes in diameter are reported for each branch order in the acute phase (3 and 7 days) and the chronic phase (14 and 21 days). Branch order of injury sites in the capillary networks of the three case examples are denoted with color coded arrows. **(C)** Representative *in vivo* image of upstream ACT vessel segments from a sham, regression, and repair event pre-injury and in the acute and chronic phase following capillary injury. Pericytes are shown in red and i.v. dye (70kDa FITC-Dextran) labeling vessels depicted in green and grayscale. Soma of ensheathing pericytes are indicated with arrows. Baseline vessel diameter is indicated (dashed blue lines) to demonstrate ACT zone constriction following capillary regression. **(D)** Graphs of percent change in the diameter of vessel segments throughout the microvascular zones (PA, ACT, Capillary, AV) of sham/repair and regression (regress.) events in animals that underwent anesthetized (blue) or awake (pink) imaging. Change from pre-injury is shown during the acute (3 or 7 days) and chronic (14 or 21 days) phase following capillary injury. ANOVA followed by Tukey’s or Dunn’s multiple comparison tests were performed depending on distribution of data. ACT zone: Acute: anesthetized sham/repair vs. regression ***p=0.0004; awake sham/repair vs. regression *p=0.014. Chronic: anesthetized sham/repair vs. regression *p=0.025; awake sham/repair vs. regression ***p=0.0006. Upstream capillary: Chronic: anesthetized regression vs. awake regression ***p=0.0002. Each datapoint is the diameter from a single vessel segment. Anesthetized: sham/repair = 10 experiments in 4 mice; regression = 9 experiments in 6 mice. Awake: sham/repair = 6 experiments in 3 mice; regression = 5 experiments in 4 mice.

We found ACT zone constriction to be a consistent phenomenon following capillary regression in the acute and chronic phase in both anesthetized and awake mice (**Fig. 3D**). In the chronic phase, capillaries upstream from regressed vessels also exhibited decreased diameters in the anesthetized state, but this was not observed in the awake state. Prominent constriction of the ACT zone was also seen as early as 10 minutes post capillary injury in awake experiments, but not in the anesthetized state (**Supp. Fig. 5**). Overall, these data demonstrate that ACT zone constriction is a lasting response irrespective of anesthetic state and occurs specifically with capillary regression after distal capillary injury.

### Constriction of the arteriole-capillary transition zone is independent of Euclidean and vessel length distance to the site of capillary injury

We next examined whether the likelihood of ACT zone constriction was related to its proximity to the injury site, and thus potential exposure to leaked blood components or inflammation. However, the Euclidean and vessel distances between the ACT zone and injury site were comparable between sham/repair and regression events (**Fig. 4A, C**). Furthermore, we did not observe a relationship between the degree of ACT zone constriction with either Euclidean or vessel distance in the acute and chronic phases (**Fig. 4B, D**). This demonstrates that proximity to the injury site does not play a role in ACT zone constriction.

**Figure 4.**
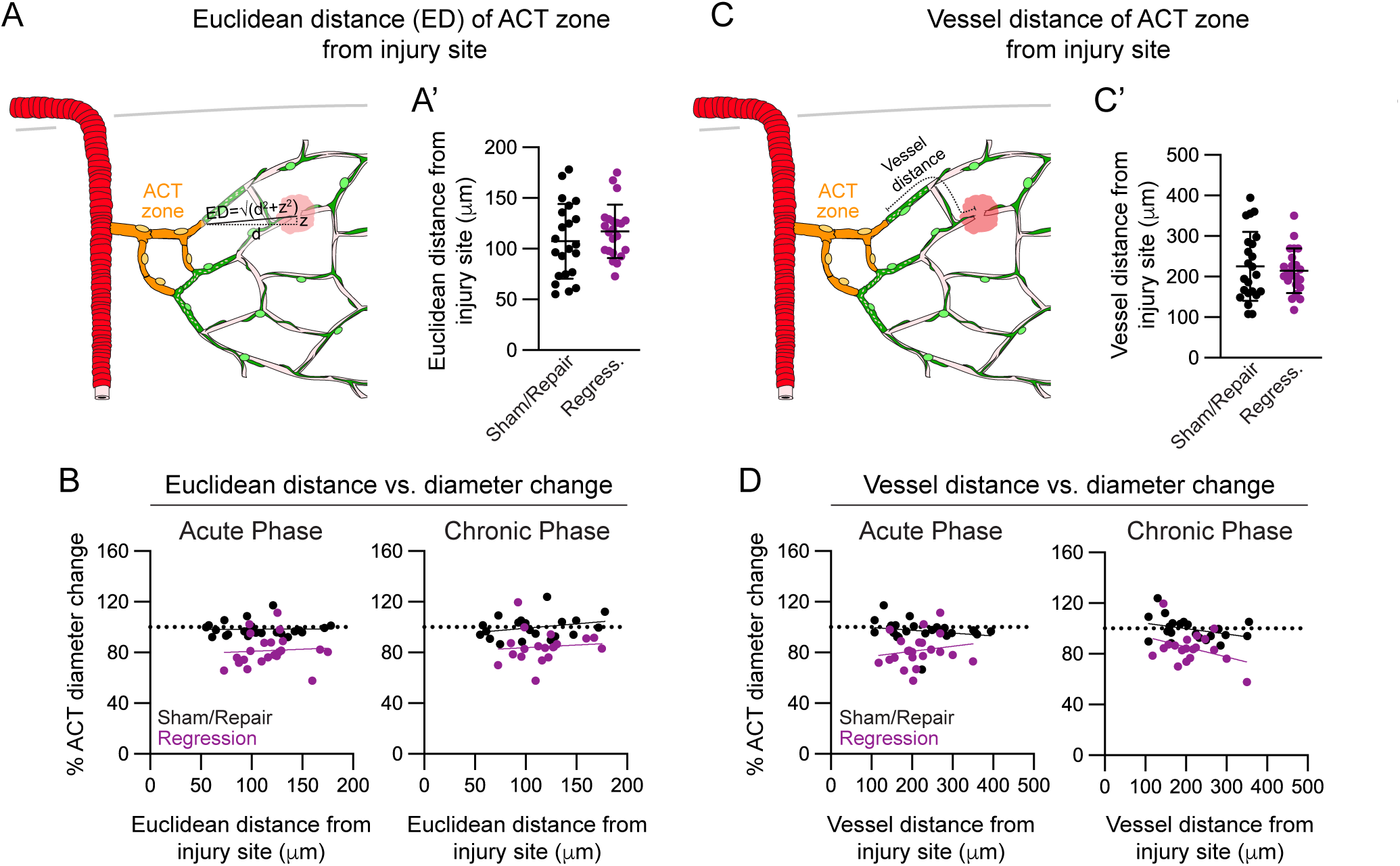
Location of capillary injury site does not influence arteriole-capillary transition zone constriction. **(A)** Schematic showing Euclidean distance between arteriole-capillary transition (ACT) zone and capillary injury site. **(A’)** Euclidean distance between ACT zone and injury site in sham/repair (black) and regression (purple) events. Unpaired t-tests revealed no significant difference. **(B)** Scatter plots of Euclidean distance of ACT zone to capillary injury site versus ACT zone diameter change from baseline during the acute (3 or 7 days) and chronic (14 or 21 days) phases. Spearman’s rank correlation tests did not show a correlation between diameter change and proximity to the injury site. (**C)** Schematic showing vessel distance between ACT zone and capillary injury site. **(C’)** Shortest vessel distance between ACT zone and injury site between sham/repair (black) and regression (purple) events. Unpaired t-tests revealed no significant difference. **(D)** Scatter plots of vessel distance versus ACT zone diameter change from baseline during the acute (3 or 7 days) and chronic (14 or 21 days) phases. Sham/repair = 16 experiments in 7 mice; regression = 14 experiments in 10 mice.

### Vasomotion of the arteriole-capillary transition zone is attenuated and uncoupled from mural cell Ca^2+^ signaling

Since mural cell Ca^2+^ signaling regulates contractile state and vessel diameter in the ACT zone^15,23^, we next investigated if resting vasomotion and mural cell Ca^2+^ dynamics were altered in the ACT zone following capillary regression. *In vivo* imaging was performed in awake PdgfrβCre-GCaMP6f mice, which express a genetically-encoded calcium sensor in mural cells. By measuring the lumen diameter and GCaMP6f fluorescence intensity of mural cells in the ACT zone (**Fig. 5A, Supp. Fig. 6A**) and penetrating arteriole (**Supp. Fig. 7A,B**) upstream of capillary injuries, we were able to study the dynamics and relationship between mural cell Ca^2+^ and vasomotion. Cross-correlation analyses were conducted to identify the timing shift needed for the strongest coupling between mural cell Ca^2+^ and vascular diameter and the correlation between these parameters, i.e., “coupling slope” (**Supp. Figs. 6B & 7C**). Diameter changes generally followed fluctuations in mural cell Ca^2+^ within 1-2 s for both the ACT zone and penetrating arterioles. Following capillary regression, we found a weaker coupling slope in the ACT zone during the acute and chronic phases (**Fig. 5B,C**) but not the penetrating arteriole (**Supp. Fig. 7D**). In contrast, sham and repair events maintained strong coupling along ACT vessels and penetrating arterioles throughout the time course (**Fig. 5C**).

**Figure 5.**
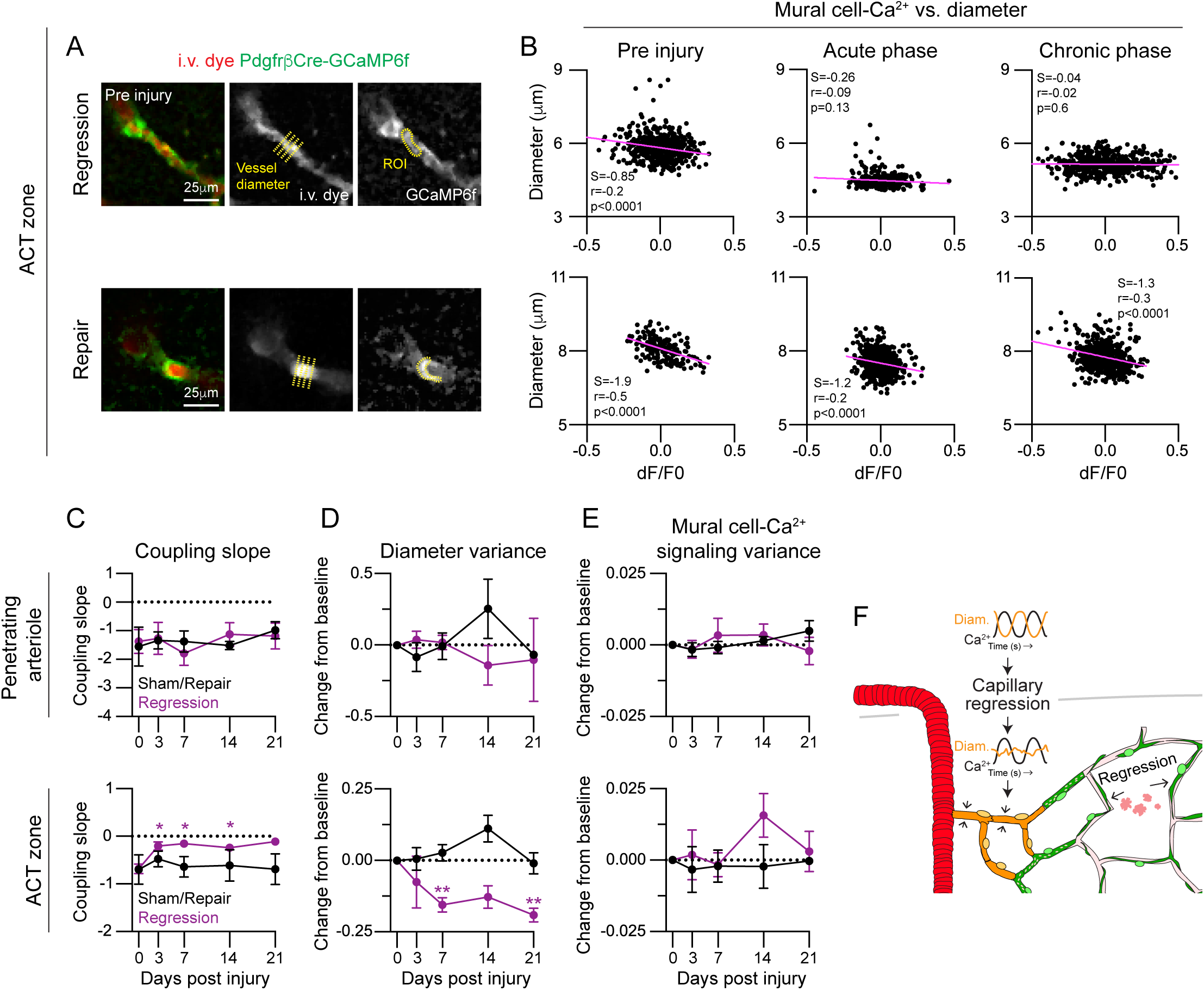
Capillary regression uncouples mural cell Ca^2+^ signaling and vessel dynamics along the arteriole-capillary transition zone. **(A)** Representative image of arteriole-capillary transition (ACT) zones from a regression and repair event in an awake PdgfrβCre-GCaMP6f mouse. Pericytes are shown in green and i.v. dye (70kDa Texas Red-Dextran) labeling vessels depicted in red. Grayscale images images show location of ACT diameter sampling (yellow crosslines) and region of interest (ROI, yellow outline) to measure GCaMP6 fluorescence intensity in ensheathing pericyte. **(B)** Scatter plots of change in mural cell GCaMP6f signal (dF/F0) versus ACT zone diameter over at least 2 minutes (data point collected every 0.512s or 1.951Hz) pre-injury, acute (3 or 7 days post-injury) and chronic (14 or 21 days post-injury). Plots are shown following cross-correlation analysis (Supplementary Fig. 6B) with strongest correlation time shift shown. Regression correlation time: Pre t=2.048s, Acute t=1.536s, Chronic t=1.024s; Repair correlation time: Pre t=1.024s, Acute t=1.024s, Chronic t=1.536s. Spearman’s rank correlations were performed, and respective r and p values are reported on graphs along with the coupling slope (S). **(C-E)** Graphs of **(C)** coupling slope, **(D)** diameter variance, and **(E)** mural cell Ca^2+^ signal variance over the course of 21 days following injury in penetrating arterioles and ACT zones upstream of sham/repair (black) and regression (purple) events. ANOVA followed by Dunnett’s multiple comparison tests were performed: Coupling slope: ACT zone - Regression events: 0 vs. 3 days: *p=0.0121, 0 vs. 7 days: *p=0.0105, 0 vs. 14 days: *p=0.0229. Diameter variance: ACT zone - Regression events: 0 vs. 7 days: **p=0.0091, 0 vs. 21 days: **p=0.0091. Sham/repair n=5, regression n=5; 4 mice. **(F)** Schematic demonstrating that capillary regression results in attenuated vasomotion (orange) and uncoupling of mural cell Ca^2+^ signaling (black) in the upstream ACT zone.

Interestingly, we found an attenuation of vasomotor oscillation (**Fig. 5D** and **Supp. Fig. 6C**), but not mural cell Ca^2+^ oscillations (**Fig. 5E** and **Supp. Fig. 6D**), in the ACT zone upstream of regressed capillaries. The mean amplitude of vessel diameter oscillations was also reduced (**Supp. Fig. 8A**), although the frequency of vasomotion of ∼0.1 Hz was maintained (**Supp. Fig. 8B**). The mean amplitude and frequency of mural cell Ca^2+^ oscillations were not affected by capillary regression (**Supp Fig. 8C, D**).

Relatively normal vascular diameter and mural cell Ca^2+^ variance was maintained along penetrating arterioles following sham, repair and regression events (**Fig. 5D, E, Supp. Fig. 7E, F**). Penetrating arterioles and ACT zones that were within the imaging field, but not directly connected to the injured vessel networks, did not exhibit any alterations to coupling slope or diameter and mural cell Ca^2+^ variance (**Supp. Fig. 9**). Altogether, these data highlight a specific decoupling between ACT zone vasodynamics and mural cell Ca^2+^ following capillary regression (**Fig. 5F**).

### Constriction of the arteriole-capillary transition zone results in broad blood flow deficits

To understand the effect of ACT zone constriction on blood flow, we used low power line-scans to track blood cell movement in both the ACT zone and uninjured, secondary off-shoots from the ACT zone in awake animals. As predicted, blood volume flux in the ACT zone (**Fig. 6A**) and blood cell flux in the secondary off-shoots (**Fig. 6B**) was reduced for at least 21 days in the event of a capillary regression, with reductions to 60-75% of basal levels. This reduction was not observed in sham injuries and repair events. Importantly, lumen diameter was unchanged in the secondary off-shoots, indicating that flow deficits were not due to local capillary constriction (**Fig. 6C**). Rather, the reduction in blood flow in the secondary off-shoots significantly correlated with the reduction in volume flux in the ACT zone (**Fig. 6D**). Thus, ACT zone constriction following capillary regression reduces blood flow into the downstream capillary bed (**Fig. 6E**).

**Figure 6.**
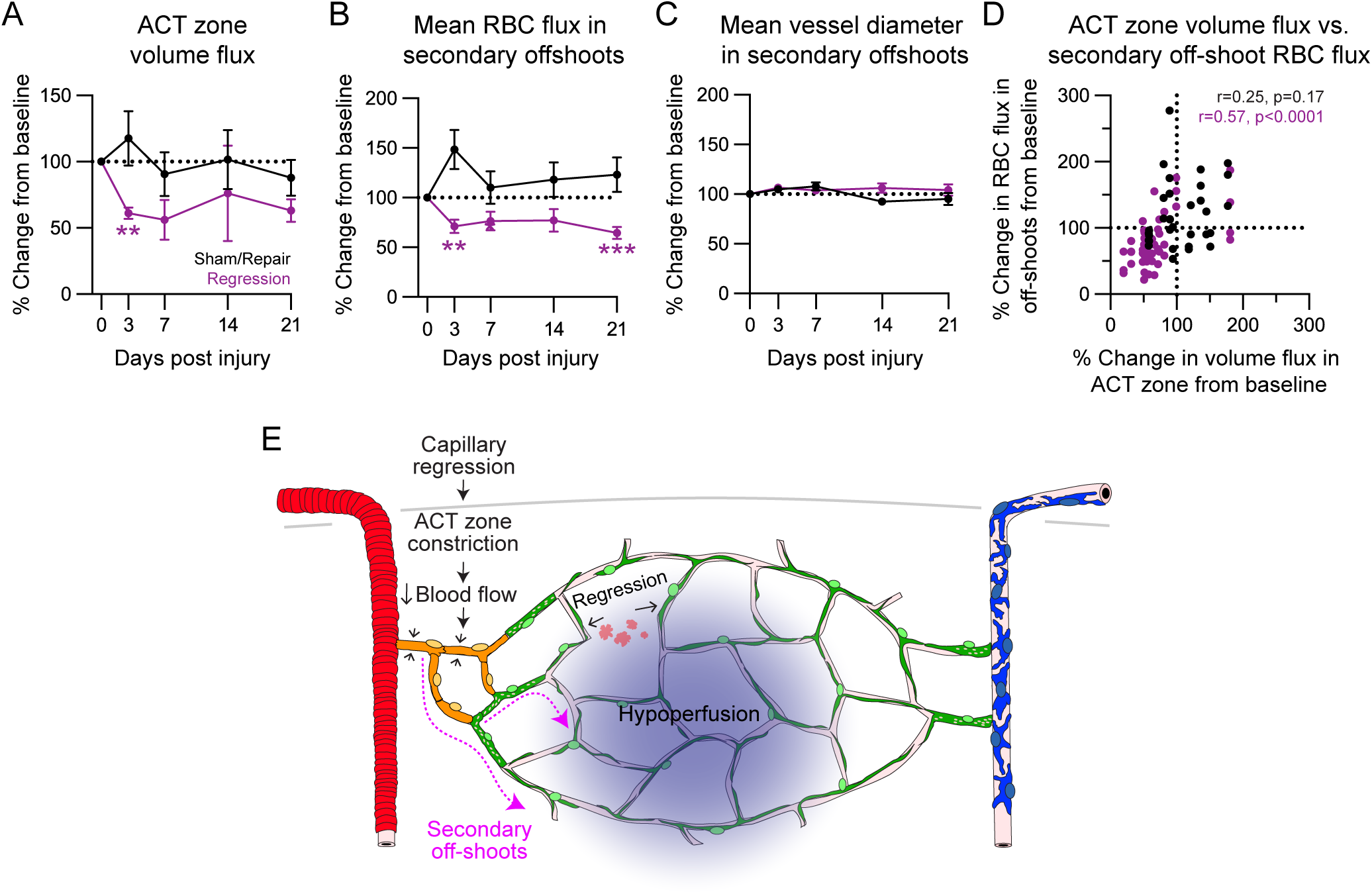
Capillary regression and chronic constriction of arteriole-capillary transition results in hypoperfusion of secondary off-shoot vessels. **(A)** Graph of upstream arteriole-capillary transition (ACT) zone blood volume flux over the course of 21 days following sham/repair (black) and regression (purple) events. ANOVA followed by Dunnett’s multiple comparison tests were performed: Regression event: 0 vs. 3 days: **p=0.0026. Sham/repair n=4, regression n=5; 4 mice. **(B, C)** Graphs of **(B)** red blood cell (RBC) flux and **(C)** vessel diameter change over the course of 21 days following injury in secondary off-shoot from ACT zone following sham/repair (black) and regression (purple) events. ANOVA followed by Dunnett’s multiple comparison tests were performed: RBC flux: Regression event: 0 vs. 3 day **p=0.0013, 0 vs. 21 days: ***p=0.0005. **(D)** Scatter plot of ACT zone blood volume flux versus RBC flux in secondary off-shoots over 21 days post-injury in sham/repair (black) and regression (purple) events. Spearman’s rank correlations were performed, respective r and p values are reported on graphs. **(E)** Schematic demonstrating that capillary regression results in ACT zone constriction reducing blood flow into the microvascular network including uninjured, secondary off-shoots.

## DISCUSSION

While the conductive nature of the brain microvasculature is involved in normal blood flow regulation, this trait may become a source of vulnerability in neurological conditions involving capillary injury. We show that injury and regression of single capillaries causes chronic and selective vasoconstriction in the upstream ACT zone, which is located hundreds of micrometers away but supplies blood to the broader capillary network. This chronic vasoconstriction reduces blood flow to 60-75% of basal levels in the ACT zone and in downstream capillaries, creating an unexpected link between capillary regression and local hypoperfusion. Capillary regression is associated with reduced amplitude of vasomotor oscillations, where mural cell Ca^2+^ dynamics are preserved but fail to translate into lumen diameter changes, suggesting a disconnection Ca^2+^ signaling and actomyosin machinery.

Prior studies have shown that the ACT zone, along with the precapillary sphincter just upstream in select arterioles, can dilate rapidly to promote blood flow during functional hyperemia^19-21,28^, or potently constrict to mediators such as endothelin-1 and ATP^29^. Relating our findings to models of neurological disease, sustained vasoconstriction has been reported after reperfusion with both focal and global cerebral ischemia^23,30^. In these prior studies, the location of constriction was most prominent in the ACT zone. Further, vasoconstriction in the ACT zone is observed with potassium-induced cortical spreading depolarization^31^, and this may contribute to lasting blood flow deficits. Note that the ACT zone has also been referred to as “precapillary arterioles” or “first/second order capillaries”, but reports of its unique sensitivity seem to be consistent despite differing nomenclature.

The preservation of GCaMP6 oscillations in mural cells suggests that Ca^2+^ handling is intact in the ACT zone after capillary regression. There are a multitude of mechanisms that could result in vasoconstriction and uncoupling of vasomotion from mural cell Ca^2+^. First, there may be impairment in the expression or function of Ca^2+^-dependent pathways that stimulate actomyosin cross-bridging (calmodulin and myosin light chain kinase), alpha-smooth muscle actin and myosin, or other cytoskeletal elements that facilitate contractility^32^. Second, it is also possible that aberrant contraction of ensheathing pericytes is Ca^2+^-independent. Rho kinase signaling promotes mural cell contraction by inhibiting myosin light chain phosphatase through rho-GTP. The upregulation of this pathway during ischemic injury and subarachnoid hemorrhage is well known^33^ and rho kinase inhibition has been show to improve blood flow by attenuating cerebral vasospasm^34^. A third possibility may be excessive vasoconstrictive tone produced by aberrant production of endothelin-1 (ET-1), a potent vasoconstrictor. ET-1 is upregulated in hemorrhagic stroke, Alzheimer’s Disease (AD), and multiple sclerosis^35^. It acts through the G-coupled protein receptor, endothelin receptor Type-A (ETAR), to induce mural cell contraction through Ca^2+^-dependent and independent mechanisms^36-38^. Finally, there may be a physical change in the vessel wall that restricts vasoreactivity, such as thickening in the vascular basement membrane^39,40^. The mechanism underlying chronic vasoconstriction in the ACT zone warrants further investigation.

We observed a difference in outcome between capillary injuries that involved regression or repair of the targeted capillary even though similar laser irradiation parameters were used (Supplementary Fig. 2). The severity of injury was comparable between groups, as both exhibited initial microbleeds followed by hyper-acute constriction within 10 minutes of injury (Supplementary Fig. 5). Acute ACT constriction may be an initial response to limit leakage and bleeding following capillary rupture. However, only those involving capillary regression were associated with chronic ACT zone vasoconstriction. It is conceivable that loss of capillary connectivity creates lasting depression or disorganization in endothelial membrane voltage and Ca^2+^ signaling, which reduces input to the ACT zone. Indeed, recent Ca^2+^ imaging studies have shown complex local Ca^2+^ signals within the endothelium, which are potentiated by conductive hyperpolarizing signals^41^. Our studies suggest that repair of the injured capillary segment, and thus re-establishment of appropriate conductive responses, is needed to alleviate ACT zone constriction.

Capillary regression is a common event in normal aging, AD, VCID, and many other progressive neurological diseases^9,10,42^. In aged animals, the ACT zone and precapillary sphincter exhibit impaired neurovascular coupling responses^43^. This has been attributed to poor responsiveness to vasomodulators. Our data suggests that there may be an additional link to capillary regression in the same vascular networks. In this regard, future studies are needed to determine whether capillary regression co-occurs with ACT zone constriction in aging and disease. Interestingly, a recent study in the aged mouse brain reported ACT *dilation* associated with age-related capillary density loss. However, cortical depth analysis revealed that deeper cortical layers exhibited more severe capillary rarefaction associated with ACT vasoconstriction^43^. Further, deep two-photon imaging in awake mice showed age-related hypoperfusion was greatest in the callosal white matter^44^. Thus, brain location may be key during studies to relate capillary regression with cerebral blood flow.

There are some limitations to these studies. First, it is unclear whether the optical ablation approach produces a form of vascular injury that is representative of disease pathology. Therefore, other targeted approaches to induce capillary regression in a zone-specific manner will be instructive in determining whether the phenomenon is technique specific. Second, we did not apply our optical ablation approach to models with disease-relevant factors, and responses in these models may differ from healthy adult mice. Third, our studies are restricted to relatively superficial layers of cortex (upper 200 μm) because optical ablations could only be effectively generated at these depths due to light scattering. Layer specific differences in vascular vulnerability and architecture will need to be considered in future studies.

In summary, our studies reveal a new dimension into how capillary-level pathology can produce outsized effects on perfusion by acting distally upon upstream ACT vessels. This link may be a contributing factor in the cerebral hypoperfusion detected in VCID, AD, and chronic phases of acute neurological injuries (stroke, hemorrhage, traumatic brain injury). Since the ACT zone is a crucial gate for blood delivery in the brain, it is imperative to understand how this vascular zone is affected in various disease conditions and whether it might be a therapeutic target for improvement of cerebral blood flow.

## MATERIALS & METHODS

### Animals

Mice were housed in a specific pathogen-free facility approved by AAALAC and were handled in accordance with protocols approved by the Seattle Children’s Research Institute IACUC committee. PdgfrβCre-tdTomato and PdgfrβCre-GCaMP6f mice were created by breeding Pdgfrβ-Cre mice (FVB and C57BL/6 × 129 background)^45^ with Ai14-flox (Jax #007914)^46^ or Ai95-flox (Jax #028865)^47^ mice (C57BL/6 backgrounds) to generate mural cell reporter lines. PdgfrβCre-tdTomato mice were crossed with Cx3cr1-GFP (Jax #005582) mice (C57BL/6 background)^48^ to simultaneously label mural cells and microglia. Both male and female mice within 3-8 months of age were utilized.

### Cranial window surgery and in vivo two-photon imaging

Chronic, skull-removed cranial windows were placed over the somatosensory cortex for *in vivo* imaging, as previously described^26^. Mice were allowed to recover for at least three weeks prior to imaging. For vascular labeling, various fluorescent dextrans were injected retro-orbitally under deep isoflurane anesthesia (2% MAC in medical-grade air). PdgfrβCre-tdTomato mice were injected with 25 µL of 5% (w/v in saline) 70kDa FITC-dextran (Sigma-Aldrich; 46945), PdgfrβCre-tdTomato; Cx3cr1-GFP mice were injected with 25 µL of 5% (w/v in saline) custom Alexa Fluor 680 (Life Technologies; A20008) conjugated to 2MDa Dextran (Fisher Scientific; NC1275021)^49^, and PdgfrβCre-Ai95 mice were injected with 25 µL of 2.5% (w/v in saline) 70kDa Texas Red^TM^ dextran (Invitrogen^TM^; D1864). During imaging of PdgfrβCre-tdTomato and PdgfrβCre-tdTomato; Cx3cr1-GFP mice, isoflurane was maintained at ∼1.5% MAC in medical-grade air. For awake imaging experiments, PdgfrβCre-Ai95 mice were habituated to head fixation on a treadmill (PhenoSys speedbelt) allowing free forward-backward movement during a 1-2 hour imaging session. For subsequent imaging experiments, mice were briefly anesthetized with isoflurane to retro-orbitally inject fluorescent dextran, and then allowed to wake up for at least 10 minutes prior to imaging. Imaging was performed with a Bruker Investigator coupled to a Spectra-Physics Insight X3. The laser was tuned for 975 nm excitation when imaging PdgfrβCre-tdTomato mice, and 920 nm excitation was used for PdgfrβCre-tdTomato; Cx3cr1-GFP and PdgfrβCre-Ai95 mice. Collection of green, red and far-red fluorescence emission was achieved with 525/70, 595/50, and 660/40 emission bandpass filters respectively, and detected with Hamamatsu GaAsP photomultiplier tubes. A 20x (1.0 NA) water-immersion objective (Olympus; XLUMPLFLN) was used to collect high-resolution images. For structural imaging of the brain vasculature, z-stacks were collected at 1.0 μm z increments at 347 μm x 347 μm (512 x 512 pixel resolution) using 3.6 μs/pixel dwell time. For blood flow analysis, three trials of 1.3 s line-scans were collected along the ACT zone (1^st^ and 2^nd^ order vessels) and uninjured, secondary capillaries with at least 10 s lags in between each line-scan. For imaging of mural cell Ca^2+^ signaling and vasomotion in PdgfrβCre-Ai95 mice, movies were captured for 307.2 s at 1.951 Hz at 295 μm x 295 μm (256 x 256 pixel resolution) averaging every two frames using 1.2 μs/pixel dwell time.

### Capillary injury using two-photon irradiation

Capillary segments to target for injuries were chosen by identifying capillary segments both within the upper 200 μm of cortex and greater than 4 branch orders from a penetrating arteriole. Injuries were induced by creating a circular line-scan path (<3 μm in diameter) to apply ∼100-154 mW of power at 800 nm excitation directly onto the vessel for 20-80 s in 20 s increments at 3.6 μs/pixel dwell time. Scanning was stopped when blood flow halted (stalled dye and/or red blood cells) and leakage of i.v. dye was observed. Sham injuries were performed in separate vascular networks using similar parameters focused on a parenchymal region adjacent to a capillary segment.

### Analysis of pericyte remodeling, vessel coverage, and pericyte-associated diameter changes

To analyze pericyte remodeling following capillary injury-induced pericyte death, remodeling processes were first identified by determining which processes of neighboring pericytes had lost pericyte contact at their terminal tips. Then, using the simple neurite tracer (SNT) plugin in FIJI, process length was measured from soma to process terminus at each imaging time point^26^. Process extension was calculated by subtracting process length at baseline from process length at each imaging time point. Vascular length uncovered by pericytes at each time point was also measured using SNT in FIJI and percent vessel coverage was based on uncovered vessel length 1 dpi. Vessel diameter was determined in pre-injury images (prior to pericyte death), the uncovered state (3 dpi) and the recovered state (14 dpi) using the VasoMetrics FIJI plugin^50^.

### Quantification of the microglia response to focal capillary injury

Maximum projected images (347 μm x 347 μm x 50 μm) were created following experiments performed in PdgfrβCre-tdTomato; Cx3cr1-GFP mice. Images encompassed the injured vessel segment and connecting vascular zones throughout all time points. Following thresholding of the Cx3cr1-GFP channel, the GFP-positive area in the 25 μm radius (1963.5 um^2^) surrounding the injury site was measured in FIJI. The percent of GFP-positive area occupying the 1963.5 um^2^ region of interest was calculated to identify the focal microglia response to capillary and sham injury. The number of Cx3cr1-GFP-positive microglia outside of the injury site in the respective images were counted using cell counter function in FIJI. Total tissue volume for each image was calculated and microglia density was normalized to μm^3^ of tissue.

### Topological analysis of vessel diameter

To analyze vessel diameter changes across the microvascular web following capillary injury and sham experiments, the territory of capillary pericytes along the vascular tree was mapped out using the SNT FIJI plugin to identify how each microvascular zone connects from penetrating arteriole to ascending venule. Experiments in which pericytes died due to capillary injury were not included. Precapillary sphincters were not easily discerned in our data sets (region of vessel narrowing with distended bulb downstream)^19^ and were therefore not separately assessed. Vessel diameter at each branch point was measured in FIJI using the VasoMetrics plugin on maximum projections of each vessel segment^50^. Vessel segments were measured at each timepoint and the percent change was calculated based on pre-injury vessel diameter.

### Analysis of distance between the injury site and the ACT zone

To measure the Euclidean distance between the injury site and the ACT zone, the distance between the focal plane of the injury site and the focal plane of the ACT zone (z) was determined. Then, the lateral distance between the injury site and the ACT zone in the x-y plane (d) was measured using the line function in FIJI on the max projected image. The equation 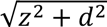 was then used to calculate the Euclidean distance. To measure the vessel distance between the injury site and the ACT zone, the length of each vessel segment between the two sites was measured using the SNT FIJI plugin.

### Analysis of vasomotion and mural cell Ca^2+^ dynamics

In awake PdgfrβCre-Ai95 mice, movies were collected of smooth muscle cells on penetrating arterioles and ensheathing pericytes on the 1^st^ and 2^nd^ order ACT zone upstream of each injury. Any imaging frames where the animal had moved, and the imaging period ∼10 s after movement, were excluded to only assess resting vasomotion. To analyze vasomotion, the VasoMetrics plugin was utilized to measure vessel diameter in each frame^50^. To analyze mural cell Ca^2+^ signatures in FIJI, images were despeckled and a ROI was drawn around the smooth muscle cells and ensheathing pericyte processes. The fluorescent intensity profile throughout the time series data was obtained using the plot z-axis profile function and the change in fluorescence divided by the median fluorescence intensity (dF/F0) was calculated. Vessel diameter and dF/F0 were plotted against each other to analyze their relationship. The coupling slope was determined by performing a cross-correlation analysis at each time point to determine the strongest inverse relationship. Raw data was used to calculate the linear regression, cross-correlation, amplitude, frequency and variance. For frequency and power analysis, MATLAB was used to bandpass filter signals from 0.025 – 0.2 Hz, the frequency range known to represent vasomotor activity^51^, and the average frequency and power were calculated. Data shown in Supplemental Figures 6A and 7B are smoothed data with a running average window of 5 data points, corresponding to 2.048 s to better show the trend of Ca^2+^ signatures and vasomotion.

### Blood flow analysis

Following collection of blood flow images using line-scans, red blood cell (RBC) velocity was measured in the ACT zone using MATLAB code for line-scanning particle image velocimetry^52^ and blood volume flux was calculated using F=⅛(π)(velocity)(diameter)^2^. In the secondary capillary offshoots, RBC flux was measured by counting the number of RBC shadows in each line-scan using the CellCounter plugin in FIJI and normalizing the numbers to represent flux over a 1 s period. Blood flow measurements from the 3 line-scans were averaged for all blood flow measurements.

### Statistics

All statistical analyses were performed in Graphpad Prism (ver. 9). Respective statistical analyses are reported in each figure legend. Normality tests, generally Shapiro-Wilk tests, were performed on necessary data sets prior to statistical tests. Standard deviation is reported in all graphs where necessary.

## ACKNOWLEDGEMENTS

SKB was supported by fellowships from the NIH/NINDS (F32NS117649) and NIH/NIA (K99AG080034). CDN was supported by a NIH training grant (T32AG052354). MJS was supported by a Diversity supplement for a grant from the NIH/NIA (R01AG062738). AYS and projects in the Shih lab were supported by grants from the NIH/NIA (R01AG062738, R21AG069375, RF1AG077731, R01AG081840). We thank G. Gürler and Liam Sullivan for comments on the manuscript.

## AUTHOR CONTRIBUTIONS

SKB, CDN, and AYS conceptualized and designed experiments. Two-photon imaging was performed by SKB and CDN. Data analysis was performed by SKB, CDN and MJS. Statistics was performed by SKB and CDN. Manuscript was written by SKB, CDN and AYS with editing and contributions from all authors.

