## Supplementary Figures for "Capillary regression leads to sustained local hypoperfusion by inducing constriction of upstream transitional vessels"

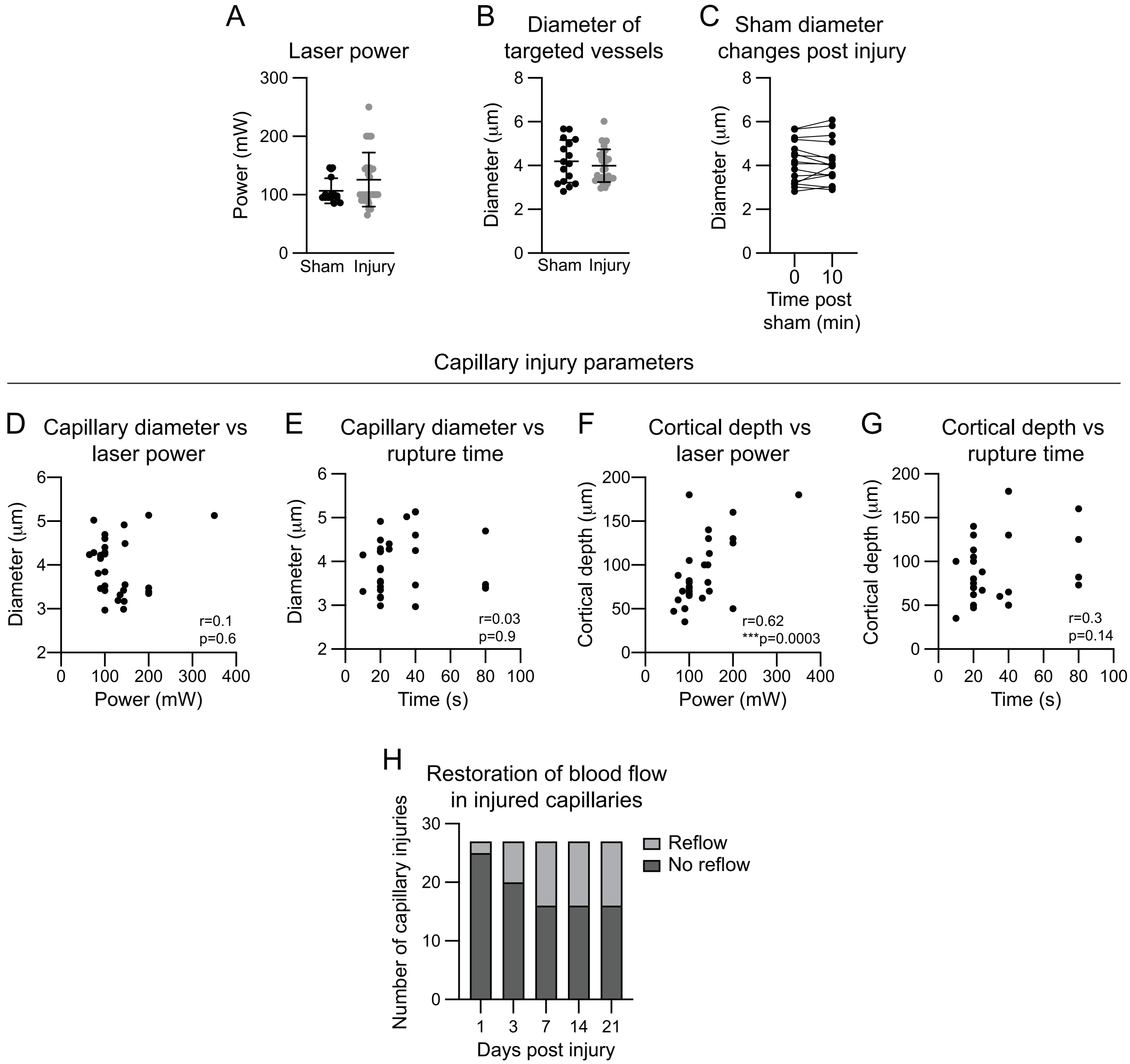


**Supplementary Figure 1. Parameters for laser-induced capillary injury.**

**(A, B)** Graphs of (A) applied laser power and (B) vessel diameter of sham (black) and capillary (gray) injuries. Appropriate parametric and non-parametric tests were used depending on distribution of data. No significant differences were detected between groups.

**(C)** Graph of vessel diameter changes for capillaries adjacent to sham injury pre and 10-minutes post injury. No significant difference detected by paired t-test.

**(D,E)** Scatter plots of vessel diameter versus (D) laser power and (E) rupture time to induce capillary injuries. Spearman's rank correlations were performed, respective r and p values are reported on graphs.

**(F,G)** Scatter plots of cortical depth versus (F) laser power and (G) rupture time to induce capillary injuries. Spearman's rank correlation test indicates increased laser power is needed to rupture capillaries deeper into the cortex.

**(H)** Graph showing the number of capillary injuries (total of 27) where blood flow was restored and reflowing (light gray) or not flowing (dark gray) for up to 21 days post capillary injury.

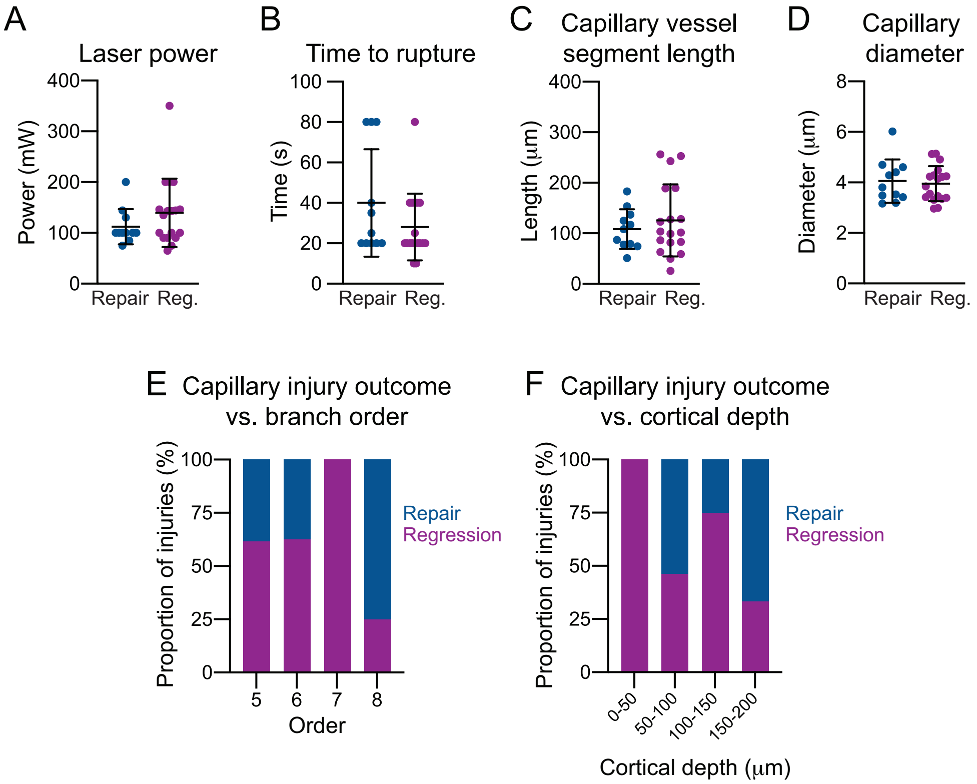


**Supplementary Figure 2. Comparison of parameters for capillary injury between regression and repair events.**
**(A-D)** Graphs of (A) applied laser power, (B) rupture time, (C) vessel segment length, and (D) vessel diameter for capillaries that repaired (blue) or regressed (Reg.; purple). Appropriate parametric and non-parametric tests were used depending on distribution of data and no significant differences were detected.
**(E,F)** Graphs showing the proportion of capillary injuries that resulted in vessel repair or regression in relation to (E) vessel branch order from the penetrating arteriole and (F) cortical depth.

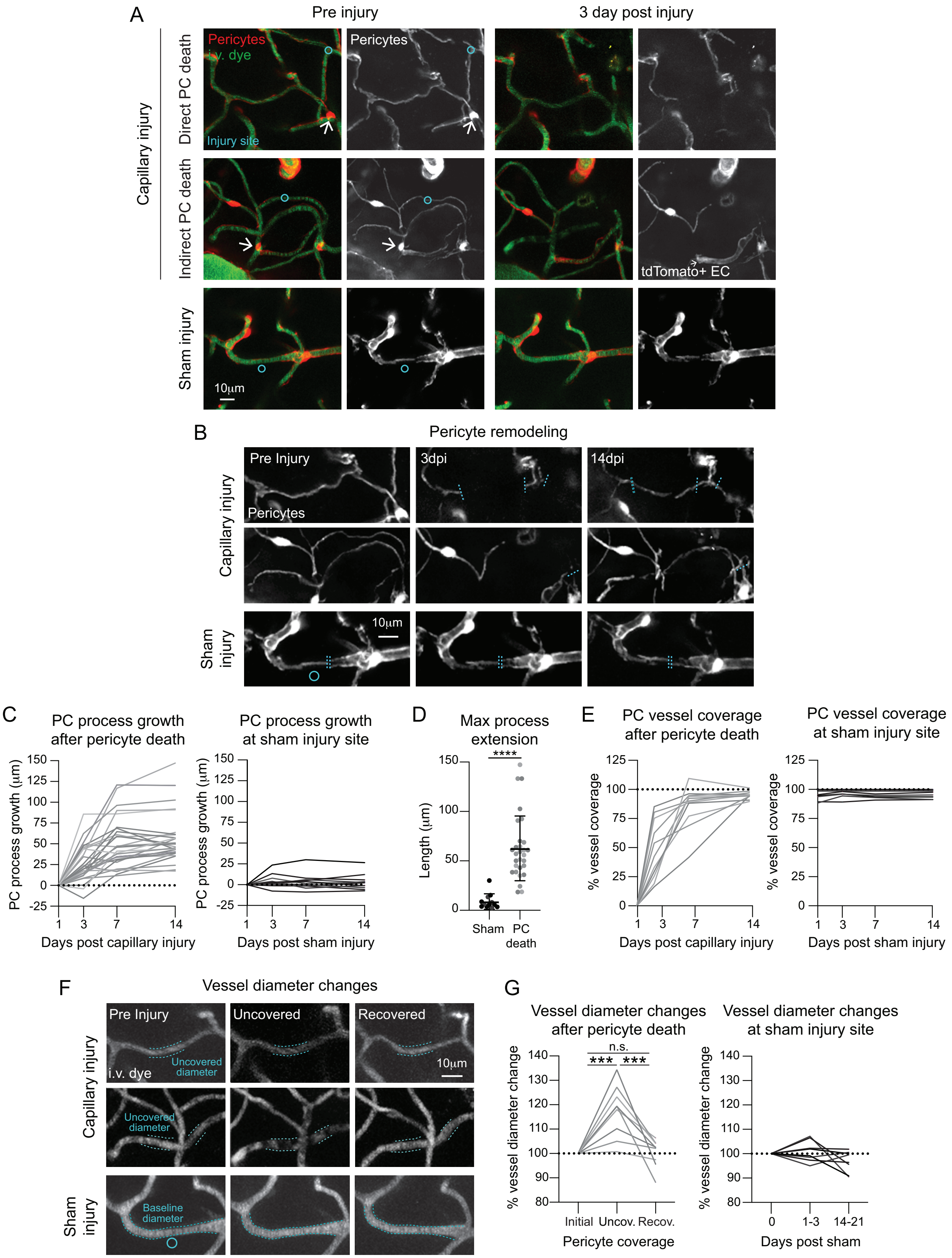
A
Pre injury
3 day post injury
Capillary injury
Direct PC death
Indirect PC death
Sham injury
Pericytes
i.v. dye
Injury site
tdTomato+ EC
10µm
B
Pericyte remodeling
Pre Injury
3dpi
14dpi
Pericytes
Capillary injury
Sham injury
10µm
C
PC process growth after pericyte death
PC process growth at sham injury site
PC process growth (µm)
Days post capillary injury
Days post sham injury
D
Max process extension
Length (µm)
****
Sham
PC death
E
PC vessel coverage after pericyte death
PC vessel coverage at sham injury site
% vessel coverage
Days post capillary injury
Days post sham injury
F
Vessel diameter changes
Pre Injury
Uncovered
Recovered
Uncovered diameter
i.v. dye
10µm
Capillary injury
Sham injury
Baseline diameter
G
Vessel diameter changes after pericyte death
Vessel diameter changes at sham injury site
% vessel diameter change
n.s.
***
***
Initial
Uncov.
Recov.
Pericyte coverage
0
1-3
14-21
Days post sham

**Supplementary Figure 3. Pericyte death may be induced by capillary injury, but remodeling of neighboring pericytes ensures coverage.**

**(A)** Representative *in vivo* images of capillary injuries (cyan circle) in a PdgfrβCre-tdTomato mouse pre and 3 days post injury that resulted in pericyte death. Direct pericyte death occurred when line-scan path injured the pericyte process (upper panel). Indirect pericyte death occurred when a nearby, uninjured pericyte died following capillary injury (middle panel). Sham injuries did not induce pericyte death (lower panel). Pericytes depicted in red and grayscale with i.v. dye (70kDa FITC-Dextran) in green. Arrows indicate pericytes that died by 3 days post injury. Note tdTomato+ endothelial cells are occasionally apparent in the PdgfrβCre-tdTomato mouse line.

**(B)** Representative *in vivo* images of pericyte remodeling following capillary injuries (upper and middle panels) and pericyte process movement following sham injuries (lower panels). Cyan dashed lines indicate the end of neighboring pericyte processes 3 days post capillary injury.

**(C)** Graphs of neighboring pericyte (PC) process growth over the course of 14 days following pericyte death (gray; left) and sham injuries (black; right). Pericyte death occurred in 7/27 of capillary injuries (in 4 mice). Sham injuries: n=8 pericyte processes, 5 mice.

**(D)** Graph of maximum process extension of pericyte processes following pericyte death and sham injuries. Mann-whitney U test detected significant differences between pericyte death and sham injuries ****$p<0.0001$.

**(E)** Graphs of pericyte (PC) vessel coverage over the course of 14 days following pericyte death (left) and sham injuries (right).

**(F)** Representative *in vivo* images of vessel diameter changes pre-injury, post pericyte death (uncovered), and when vessels were recovered by pericyte processes following capillary (upper 2 panels) and sham (lower panel) injuries. Dashed lines outline uncovered vessel diameter size in uncovered state for capillary injuries and baseline diameter for sham injuries.

**(G)** Graphs of vessel diameter changes during pre-injury, uncovered, and recovered states for capillary injuries that resulted in pericyte death (left). Sham injuries (right) show diameter changes at 1-3 and 14-21-days post sham. ANOVA followed by Tukey's multiple comparison tests were performed: pericyte death: initial vs. uncovered ***$p=0.0002$, uncovered vs. recovered ***$p=0.0003$.

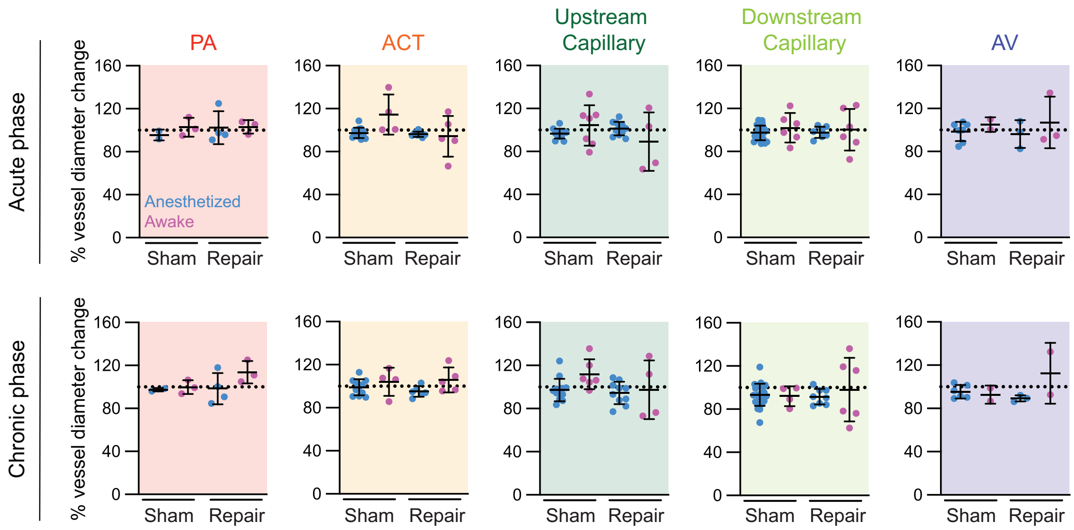


**Supplementary Figure 4. Comparison of vessel diameter changes along the vascular zones following repair and sham injury events.**

Graphs of percent change in the diameter of vessel segments throughout the microvascular zones encompassing penetrating arterioles (PA), arteriole-capillary transition zones (ACT), capillaries, and ascending venules (AV) of sham and repair events in animals that underwent anesthetized (blue) or awake (pink) imaging. Change from pre-injury is shown during the acute (3 or 7 days) and chronic (14 or 21 days) phase following capillary or sham injury. ANOVA tests were performed, and no significant differences were detected. Each datapoint is the diameter from a single vessel segment. Anesthetized: sham = 6 experiments in 4 mice; repair = 4 experiments in 4 mice; Awake: sham = 3 experiments in 3 mice; repair = 3 experiments in 3 mice.

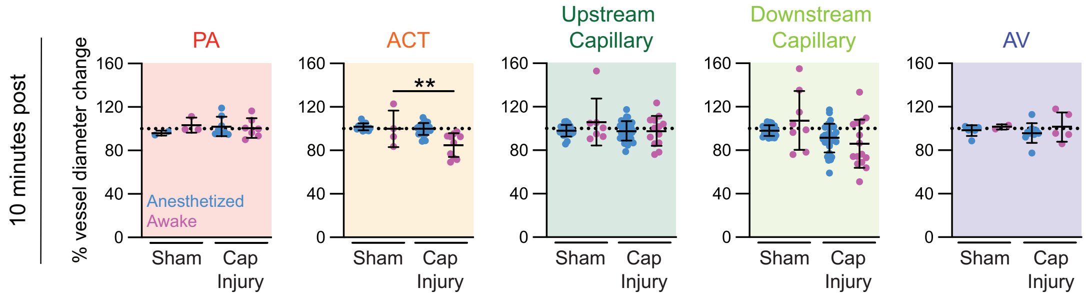


**Supplementary Figure 5. Arteriole-capillary transition vessels constrict as early as 10 minutes post-optical capillary injury in awake animals.**

Graphs of percent change in the diameter of vessel segments 10 minutes post capillary injury or sham throughout the microvascular zones, encompassing penetrating arterioles (PA), arteriole-capillary transition zones (ACT), capillaries, and ascending venules (AV) of sham and repair events in animals that underwent anesthetized (blue) or awake (pink) imaging. Change from pre-injury is shown during the acute (3 or 7 days) and chronic (14 or 21 days) phase following capillary or sham injury. ANOVA followed by Tukey's or Dunn's multiple comparison tests were performed depending on distribution. ACT zone: Awake sham vs. capillary injury **p=0.0096. Each datapoint is the diameter from a single vessel segment. Anesthetized: sham = 6 experiments in 4 mice; capillary injuries = 14 experiments in 7 mice. Awake: sham = 3 experiments in 3 mice; capillary injury = 8 experiments in 4 mice.

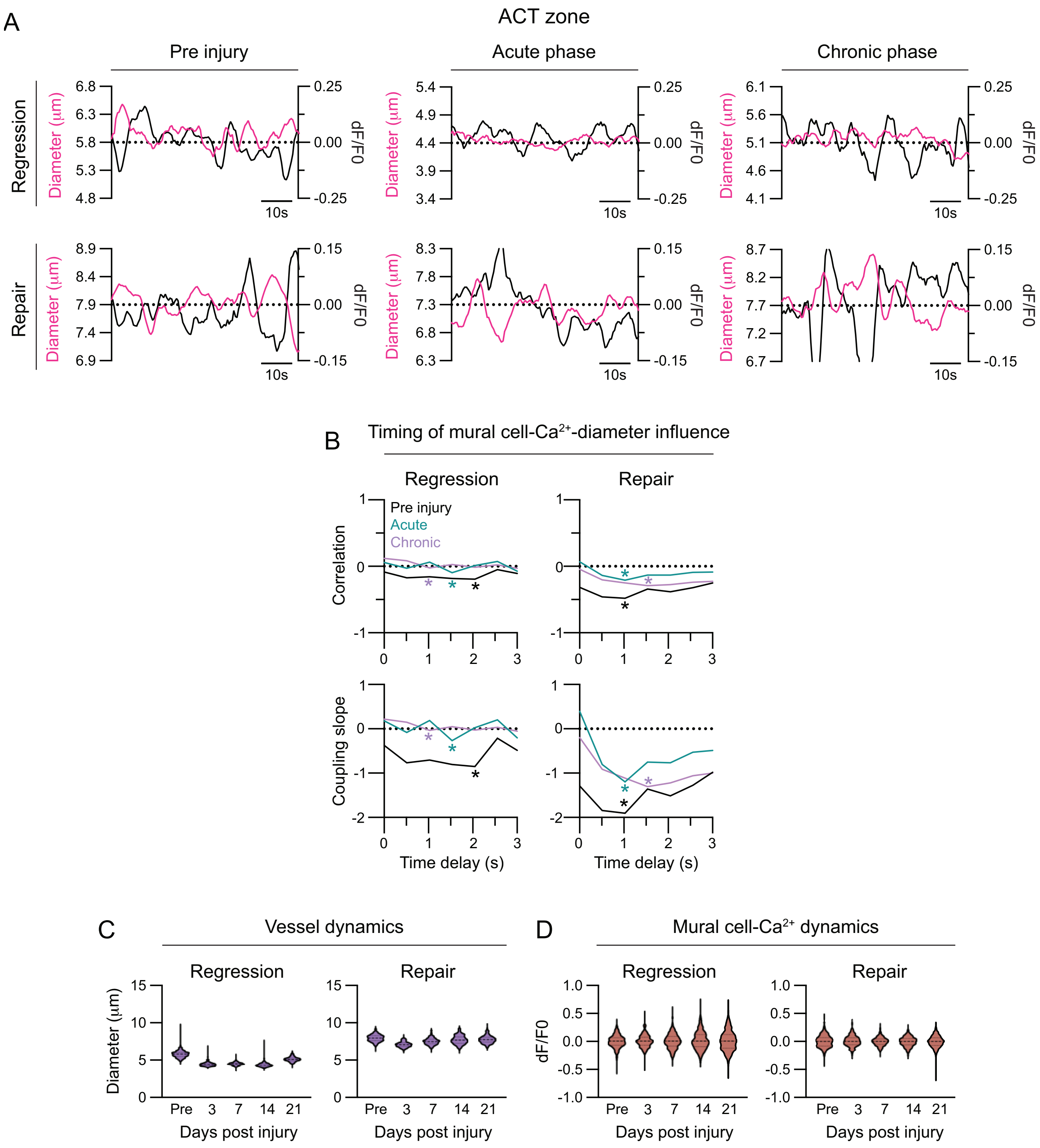


**Supplemental Figure 6. Mural cell-calcium and vessel diameter dynamics in arteriole-capillary transition zone following capillary injury.**
**(A)** Graphs showing case examples of change in arteriole-capillary transition (ACT) zone vessel diameter and mural cell-$Ca^{2+}$ signaling (dF/F0) over 1 minute prior to injury and during the acute (3 or 7 days) and chronic (14 or 21 days) in a capillary regression and repair event.

**(B)** Graphs showing case examples of cross correlation analysis and respective coupling slopes within 3 seconds of ACT zone vessel diameter and mural cell-$Ca^{2+}$ signaling changes for a capillary regression and repair event. Strongest influence of mural cell-GCaMP6f signaling on ACT vasodynamics are indicated with asterisks and based on the strongest correlation for experiments pre-injury (black) and in the acute (green) and chronic (purple) phase.
**(C, D)** Graphs showing the distribution of (C) vessel diameter and (D) mural cell-$Ca^{2+}$ signaling (dF/F0) in the ACT zone over at least 2 minutes prior to injury and 3-, 7-, 14-, 21-days post injury in a regression and repair event.

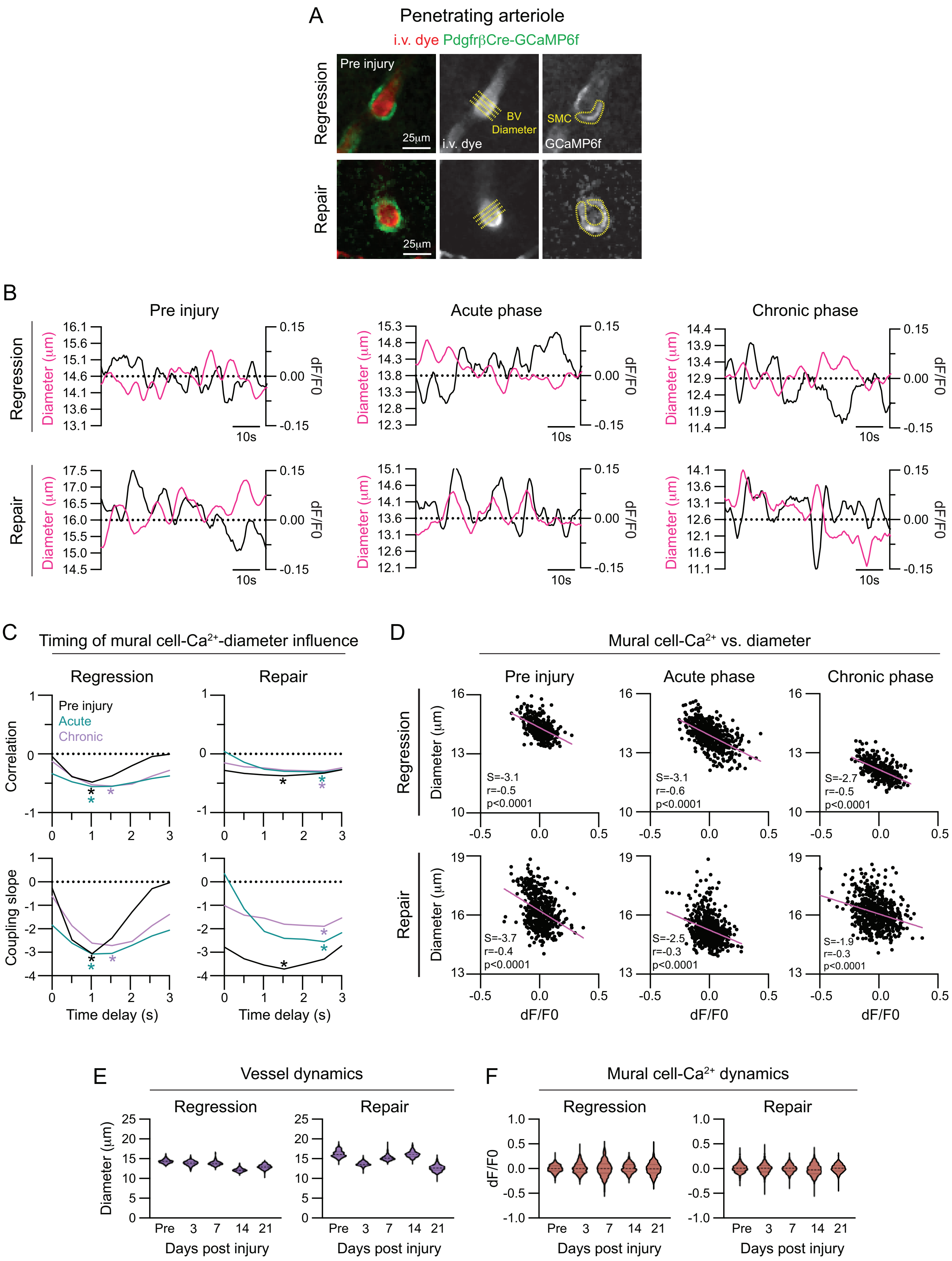
A
Penetrating arteriole
i.v. dye PdgfrβCre-GCaMP6f
Regression
Repair
Pre injury
25μm
BV
Diameter
SMC
i.v. dye
GCaMP6f
B
Pre injury
Acute phase
Chronic phase
Diameter (μm)
dF/F0
10s
C
Timing of mural cell-Ca2+-diameter influence
Regression
Repair
Pre injury
Acute
Chronic
Correlation
Coupling slope
Time delay (s)
D
Mural cell-Ca2+ vs. diameter
Pre injury
Acute phase
Chronic phase
S=-3.1
r=-0.5
p<0.0001
S=-3.1
r=-0.6
p<0.0001
S=-2.7
r=-0.5
p<0.0001
S=-3.7
r=-0.4
p<0.0001
S=-2.5
r=-0.3
p<0.0001
S=-1.9
r=-0.3
p<0.0001
dF/F0
E
Vessel dynamics
Days post injury
Pre 3 7 14 21
F
Mural cell-Ca2+ dynamics
dF/F0

**Supplemental Figure 7. Mural cell-calcium and vessel diameter dynamics are not altered along penetrating arterioles following capillary injury.**

**(A)** Representative *in vivo* t-series image of penetrating arteriole (PA) from a regression and repair event in an awake PdgfrβCre-GCamP6f mouse. Smooth muscle cells (SMCs) are shown in green and i.v. dye (70kDa Texas Red-Dextran) labeling vessels depicted in red. In respective grayscale images, an example of vessel diameter (yellow crosslines) and GCaMP6 fluorescent intensity (yellow outline) analysis is shown.

**(B)** Graphs showing case examples of change in PA vessel diameter and mural cell-$Ca^{2+}$ signaling (dF/F0) over 1 minute prior to injury and during the acute (3 or 7 days) and chronic (14 or 21 days) phases in a capillary regression and repair event.

**(C)** Graphs showing case examples of cross correlation analysis and respective coupling slopes within 3 seconds of PA vessel diameter and mural cell-$Ca^{2+}$ signaling changes in a capillary regression and repair event. Strongest influence of mural cell-$Ca^{2+}$ signaling on PA vasodynamics are indicated with asterisks and based on the strongest correlation for experiments pre-injury (black) and in the acute (green) and chronic (purple) phase.

**(D)** Scatter plots of change in mural cell-$Ca^{2+}$ signal (dF/F0) versus PA diameter over at least 2 minutes (data point collected every 0.512s or 1.951Hz) pre-injury, acute (3 or 7 days post injury) and chronic (14 or 21 days post injury) phase. Plots are shown following cross-correlation analysis (Supplementary Fig. 7C) with strongest correlation. Regression correlation time: Pre t=1.024s, Acute t=1.024s, Chronic t=1.536s; Repair correlation time: Pre t=1.536s, Acute t=2.56s, Chronic t=2.56s. Spearman's rank correlations were performed, respective r and p values are reported on graphs along with the coupling slope (S).

**(E,F)** Graphs showing distribution of (E) vessel diameter and (F) mural cell-$Ca^{2+}$ signaling (dF/F0) in the PA over at least 2 minutes prior to injury and 3-, 7-, 14-, 21-days post-injury in a regression and repair event.

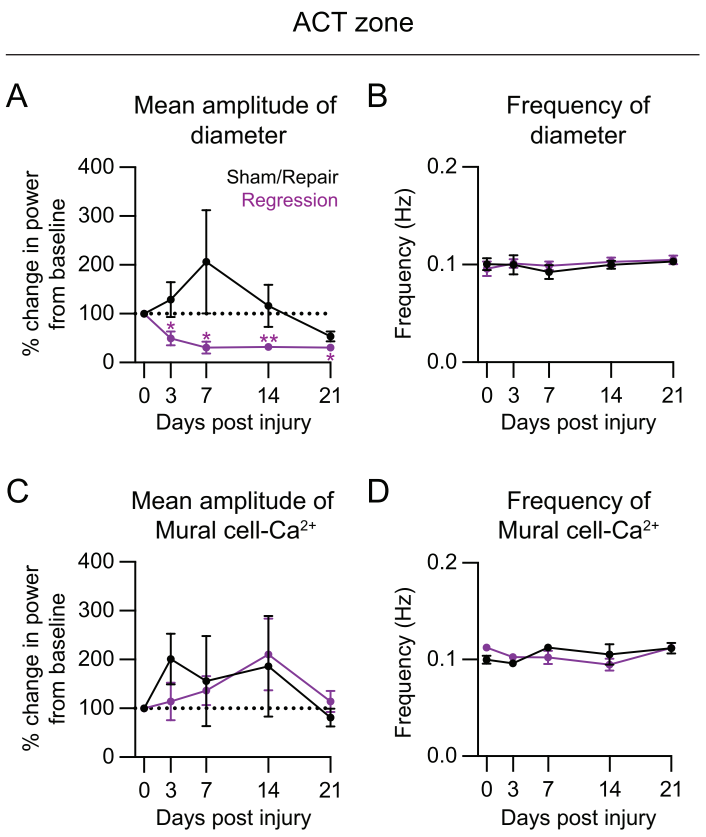


**Supplemental Figure 8. Power and frequency of mural cell-calcium is maintained in the arteriole-capillary transition zone.**

**(A & C)** Graphs showing percent (%) change in mean amplitude of (A) vessel diameter and (C) mural cell-$Ca^{2+}$ signaling in the arteriole-capillary transition (ACT) zone from baseline over the course of 21 days upstream of sham/repair (black) and regression (purple) events. ANOVA followed by Dunnett's multiple comparison tests were performed: Mean diameter amplitude- Regression event: 0 vs. 7 days: *p=0.0435, 0 vs. 14 days: **p=0.0018, 0 vs 21 days: *p=0.0122. Sham/repair n=5, regression n=5; 4 mice.

**(B & D)** Graphs showing mean frequency of (B) ACT diameter and (D) mural cell-$Ca^{2+}$ signaling over the course of 21 days upstream of sham/repair and regression events. ANOVA tests detected no significant differences in frequency of diameter or mural cell-$Ca^{2+}$ signaling.

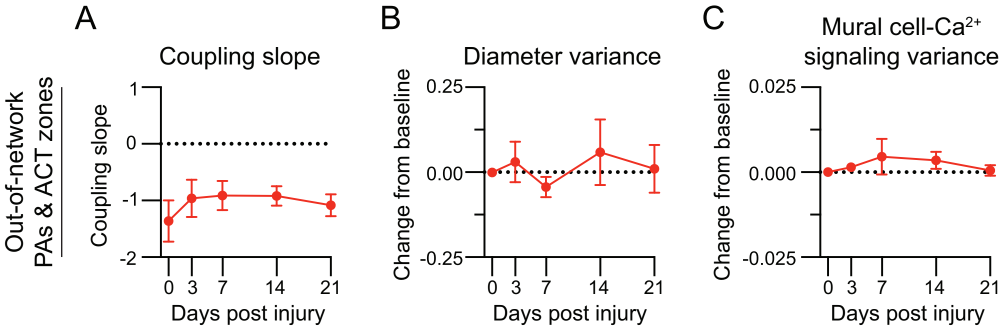


**Supplemental Figure 9. Mural cell-calcium and vessel diameter dynamics are not altered in out-of-network penetrating arterioles and arteriole-capillary transition zones.**
**(A-C)** Graphs of (A) coupling slope, (B) change in diameter variance, and (C) change in mural cell-$Ca^{2+}$ variance over the course of 21 days in out-of-network penetrating arterioles (PAs) and arteriole-capillary transition (ACT) zones. Vessels were within imaging frames but not a part of the injured microvascular networks. ANOVA tests detected no significant differences. n=5 vessels; 4 mice.
